# The female pheromone (*Z*)-4-undecenal mediates flight attraction and courtship in *Drosophila melanogaster*

**DOI:** 10.1101/2021.01.06.425638

**Authors:** Felipe Borrero-Echeverry, Marit Solum, Federica Trona, Erika A. Wallin, Marie Bengtsson, Peter Witzgall, Sebastien Lebreton

## Abstract

Specific mate communication and recognition underlies reproduction and hence speciation. Mate communication evolves during adaptation to ecological niches and makes use of social signals and habitat cues.

Our study provides new insights in *Drosophila melanogaster* premating olfactory communication, showing that female pheromone (*Z*)-4-undecenal (*Z*4-11Al) and male pheromone cVA interact with food odour in a sex-specific manner. Furthermore, *Z*4-11Al, which mediates upwind flight attraction in both sexes, also elicits courtship in experienced males.

Twin variants of the olfactory receptor Or69a are co-expressed in the same olfactory sensory neurons, and feed into the same glomerulus in the antennal lobe. *Z*4-11Al is perceived via Or69aB, while the food odorant (*R*)-linalool is a main ligand for the other variant, Or69aA. That *Z*4-11Al mediates courtship in experienced males, not *(R)-* linalool, is probably due to courtship learning. Behavioural discrimination is reflected by calcium imaging of the antennal lobe, showing distinct glomerular activation patterns by these two compounds.

Male sex pheromone cVA is known to affect male and female courtship at close range, but does not elicit upwind flight attraction as a single compound, in to contrast to *Z*4-11Al. A blend of cVA and the food odour vinegar attracted females, while a blend of female pheromone *Z*4-11Al and vinegar attracted males instead.

Sex-specific upwind flight attraction to blends of food volatiles and male and female pheromone, respectively, adds a new element to *Drosophila* olfactory premating communication and is an unambiguous paradigm for identifying the behaviourally active components, towards a more complete concept of food-pheromone odour objects.

**Summary:** The female-produced, species-specific volatile pheromone of *D. melanogaster* attracts both sexes from a distance, alone and in concert with food odorants, and elicits courtship in experienced males.

## Introduction

Structure and function of the olfactory system have been studied in *Drosophila* like in no other insect, from peripheral odorant perception to central pathways generating behavioural output. And, a particular emphasis of this work has been placed on courtship, inspired by a characteristic behavioural response (Depetris-Chauvin et al. 2015, Kohl et al. 2015, Auer and Benton 2016, Grabe et al. 2016, Bates et al. 2020). The combined molecular and physiological know-how enables even investigations of the evolutionary development of olfactory channels in response to pheromones (Khallaf et al. 2020a,b) and host odorants (Dekker et al. 2006, Markow 2019, Auer et al. 2020), across *Drosophila* phylogenies.

What is still unclear is whether olfactory tuning and response to pheromones and host or food odours evolve independently. And, although odorants enable evaluation and recognition of the emitter or source from a distance, little attention has been paid to the role of species-specific sex pheromones in distant mate recognition and flight attraction, prior to courtship enactment.

Drosophila cuticular hydrocarbon (CHC) profiles are sexually dimorphic, they vary and rapidly diverge between species and therefore contribute to sexual, reproductive isolation (Howard et al. 2003, Legendre et al. 2008, Alves et al. 2010, De Oliveira et al. 2011, Davis et al. 2020). Naturally, this congruently applies to their volatile oxidation products, particularly monounsaturated aldehydes, which derive from diunsaturated CHCs.

A CHC produced by female flies, (*Z*,*Z*)-7,11-heptacosadiene (7,11-HD), has been shown to afford reproductive isolation between *D*. *melanogaster* and its sibling species *D*. *simulans.* It is perceived by gustatory receptors owing to its low volatility (Billeter et al. 2009, Thistle et al. 2012, Toda et al. 2012, Billeter and Wolfner 2018, Seeholzer et al. 2018, Sato and Yamamoto 2020). Oxidation of 7,11-HD gives rise to the volatile pheromone (*Z*)-4-undecenal (*Z*4-11Al) which is perceived by one of the two variants of the olfactory receptor DmelOr69a (Or69a) and elicits flight attraction in males and females (Lebreton et al. 2017). The Or69a gene encodes two or more proteins, in a range of *Drosophila* species (Robertson et al. 2003, Conceicao and Aguade 2008). In *D*. *melanogaster*, the two isoforms of Or69a, Or69aA and Or69aB, are tuned to food odorants, including (R)-linalool and to the female pheromone *Z*4-11Al, respectively. They are expressed in the same olfactory sensory neurons (OSNs), and afford combined input of sex and food stimuli (Lebreton et al. 2017).

In comparison, *cis*-vaccenyl acetate (cVA) is a pheromone produced by males that increases female receptivity and reduces male attraction to recently mated females (Bartelt et al. 1985, Ejima et al. 2007, Kurtovic et al. 2007, Keleman et al. 2012, Lebreton et al. 2014). However, cVA cannot account for specific mate communication since it is shared by many other *Drosophila* species (El-Sayed 2020). The processing of cVA stimuli in sexually dimorphic neural pathways has been mapped from antennal input to third-order neurons in the lateral horn, where gender-specific courtship behaviour is generated (Kohl et al. 2013, Clowney et al. 2015).

cVA activates the sexually dimorphic fruitless *(Fru)* circuitry that controls male courtship (Manoli et al. 2005, Stockinger et al. 2005, Billeter et al. 2006, Cachero et al. 2010), whereas *Z*4-11Al is not part of this circuit. Since *Z*4-11Al is produced by females, we investigated its effect on flight attraction and courtship, which are successive steps in male reproductive behaviour. An upwind flight assay confirms that *Z*4-11Al elicits attraction in naive males (Lebreton et al. 2017), while it elicits courtship only in experienced, previously mated males.

Remarkably, males distinguished between *Z*4-11Al and the food odorant (*R*)-linalool, which did not elicit courtship. Both compounds are perceived via the two isoforms of Or69a, that are co-expressed in the same OSNs, while functional imaging of the antennal lobe (AL) showed nonetheless different activation patterns for *Z*4-11Al and (*R*)-linalool.

Food odours act as aphrodisiacs and promote courtship in *Drosophila* (Grosjean et al. 2011, Gorter et al. 2016, Ando et al. 2020). Vinegar has traditionally been used as a source of food odorants in fruit fly research, and has a prominent synergistic effect on cVA perception and attraction (Lebreton et al. 2015, Das et al. 2017, Cazale-Debat et al. 2019). Vinegar headspace contains several active compounds (Becher et al. 2010) but greatly varies in composition between makes and types (Callejón et al. 2009, Chinnici et al. 2009). Over-ripe fermenting fruit, on the other hand, is a feeding, mating, and oviposition site for fruit flies. Yeasts growing on fruit release a very rich volatome, that accounts for strong fly attraction (Becher et al. 2012, Buser et al. 2014, Christiaens et al. 2014, Ljunggren et al. 2019).

We compared the effect of vinegar and yeast headspace on sex attraction to pheromone. Blends of vinegar odour and male or female pheromone produced sex-specific flight attraction of females and males, respectively. In comparison, both sexes respond to blends of yeast and cVA or *Z*4-11Al.

## Materials and methods

### Insects and chemicals

Flies were reared on a standard sugar-yeast-cornmeal diet at room temperature (19 to 22°C) under a 16:8-h L:D photoperiod. Newly eclosing flies were anesthetized with CO2 and sexed under a microscope. Virgin flies were identified by the presence of meconium, and were kept together with flies of the same sex. Mated males were kept together with females, and were separated 2 d before experiments. Flies were kept in 30-mL Plexiglas vials with fresh food, and were six d old when used for experiments.

Isomeric purity of (*Z*)-4-undecenal (Z4-11Al) and (*E*)-4-undecenal (E4-11Al) was 98.6% and 97.8%, repectively, according to gas chromatography coupled to mass spectrometry (6890 GC and 5975 MS, Agilent Technologies, Santa Clara, CA, USA). Chemical purity of the synthetic aldehydes was >99.9%, and of the other test chemicals >97%. Heptane (redistilled; Merck, Darmstadt, Germany) was used as solvent.

### Functional imaging

The head capsule was opened by incising the cuticle between the antennae and the eyes. With the brain immersed in Ringer’s saline, the ALs were exposed by removing muscle tissue, glands and trachea. In vivo recordings of illuminated preparations were processed using custom software (Strutz et al. 2014).

We used a Till Photonic imaging system with an upright Olympus microscope (BX51WI) and a 20x Olympus objective (XLUM Plan FL 20x/0.95W). A Polychrome V provided light excitation (475 nm), which was then filtered (excitation: SP500, dicroic: DCLP490, emission LP515) and captured by a CCD camera (Sensicam QE, PCO AG) with symmetrical binning. For each measurement, a series of 300 frames was taken (1 Hz). Data were analyzed using IDL (Research Systems Inc., Boulder, CO, USA).

A 3-D map of the fruit fly AL (Grabe et al. 2015) served to link the active area to individual glomeruli. All experimental flies contained the calcium dependent fluorescent sensor G-CaMP 3.0 (Nakai et al. 2001) together with a promoter Gal4 insertion to direct expression of the calcium sensor to specific neuron populations. Stimulus-evoked fluorescence in these flies arises from the population of labeled neurons that are sensitive to the specific odour.

We tested the physiological responses in input neurons, i.e. the axonal terminals of OSNs in the AL. Mass labelling of olfactory sensory neurons (OSNs) was achieved using the transgenic line Orco-GAL4 that drives expression in at least 60% of all OSNs (Larsson et al. 2004). Additionally, for mass labelling of PNs, the transgenic line GH146-GAL4, expressing G-CaMP in 83 out of the 150 PNs in each AL (Vosshall and Stocker 2007) has been used. Fly lines were obtained from the Bloomington *Drosophila* Stock Center (Indiana University, Bloomington, IN, USA). Transgenic flies have been generated using standard procedures.

### Courtship assay

Wild-type flies for courtship experiments were the *D. melanogaster* strains Dalby-HL (Ruebenbauer et al. 2008) and Canton-S, and and the sister species *D. simulans.*

Orco-Gal4/uas-Or69aRNAi flies, and the Orco-Gal4 and uas-Or69aRNAi (VDCR, Vienna) parental lines were used to confirm the effect of *Z*4-11Al on courtship behavior. Canton-S served as comparison for knockouts of the same background. We resorted to the Orco-Gal4 line, since males of the Or69a-Gal4 line did not court. Because of the broad expression of the Or69aRNAi, we cannot exclude that other Ors involved in courtship in experienced males are also affected.

Courtship experiments were done on a light box (37×28×2.5 cm; color temperature 5.000 ± 5% Kelvin; Kaiser Slimlite 2422, Kaiser Fototechnik GmbH & Co. KG, Buchen, Germany). The courtship arena consisted of three glass plates (17×13×0.5 cm) placed on top of each other. Twelve circular holes (ø 3 cm) were cut in the middle plate to form 12 circular single-pair mating chambers. All glassware was heated to 350°C for 8 h before use. Tests were done between 2 and 5 h after onset of scotophase (16:8 L:D photoperiod). Target flies, unmated males or females, were anesthetized on ice and decapitated before experiments. Test flies were either mated or unmated males. A single target fly was added to each mating chamber, and 1 μl heptane (control) or 1 μl heptane containing 1 ng *Z*4-11Al was applied onto its abdomen. After solvent evaporation (2 min), a single live test fly was introduced into each mating chamber and observed during 20 min. Males were scored when showing vigorous courtship behaviour, including wing vibration, licking, and attempted copulation. Treatments (*n* = 80) and controls (*n* = 80) were conducted simultaneously, and each fly was tested once.

### Food odour collection

Brewer’s yeast, *Saccharomyces cerevisiae* (strain S288C), was grown in minimal media (Merico et al. 2007) during 20 h in a shaking incubator at 25°C and 260 RPM. 50 mL yeast broth or white vine vinegar (7.1%, Zeta, Sweden) were filled into a wash bottle and charcoal filtered air was bubbled through the bottle at a rate of 200 mL/min. Headspace was trapped in 4×40-mm glass tubes holding 300 mg of Porapak Q 50/30 (Waters Corporation, Milford, Ma, USA) during 2 h (Becher et al. 2012). These air filters were eluted with 1.2 mL redistilled ethanol (Labscan), the eluate was sprayed during 2 h of wind tunnel experiments.

### Wind tunnel assay

Upwind flight attraction was observed in a glass wind tunnel (30×30×100 cm). The wind tunnel was lit diffusely from above, at 13 lux, temperature ranged from 20°C to 24°C, relative humidity from 38% to 48% and charcoal filtered air (Camfil AB, Malmö, Sweden), at a velocity of 0.25 m/s, was produced by a fan (Fischbach GmbH, Neunkirchen, Germany). Yeast headspace and odour blends were delivered from the centre of the upwind end of the wind tunnel via a piezo-electric microsprayer (Becher et al. 2010).

Forty flies were flown individually to each treatment. Flies were scored when flying upwind over 80 cm in the wind tunnel centre, from the release cage towards the odour source. Unmated, fed, 3-d-old males and females were flown towards yeast and vinegar headspace alone and in blends with *Z*4-11Al sprayed at 10 ng/min (Lebreton et al. 2017) and 300 ng/min *cis*-vaccenyl acetate (cVA) (Lebreton et al. 2015). Isomeric purity of the synthetic compounds, according to gas chromatography, was 98.6%. and 97.3%, respectively.

### Statistical analysis

Generalized linear models (GLM) with a Bernoulli binomial distribution were used to analyse wind tunnel data. Landing at source and sex were used as the target effects. *Post-hoc* Wald pairwise comparison tests were used to identify differences between treatments. Courtship assays were analysed using chi-squared tests to compare between treatments and their respective control. Statistical analysis was calculated with R (R Core Team 2013) and SPSS Version 22 (IBM Corp.).

## Results

### Flight attraction to blends of pheromone and food odour

The male-produced pheromone cVA (Bartelt et al. 1985) is a core paradigm for behavioural and neurophysiological studies in *Drosophila.* The volatile, female-produced and species-specific pheromone *Z*4-11Al has been discovered more recently (Lebreton et al. 2017). We therefore compared flight attraction to male pheromone cVA and female pheromone *Z*4-11Al, in *D*. *melanogaster* males and females. Pheromones were tested alone and in blends with odorants from vinegar and yeast, signalling food sources (Figure 1).

**Figure 1.**
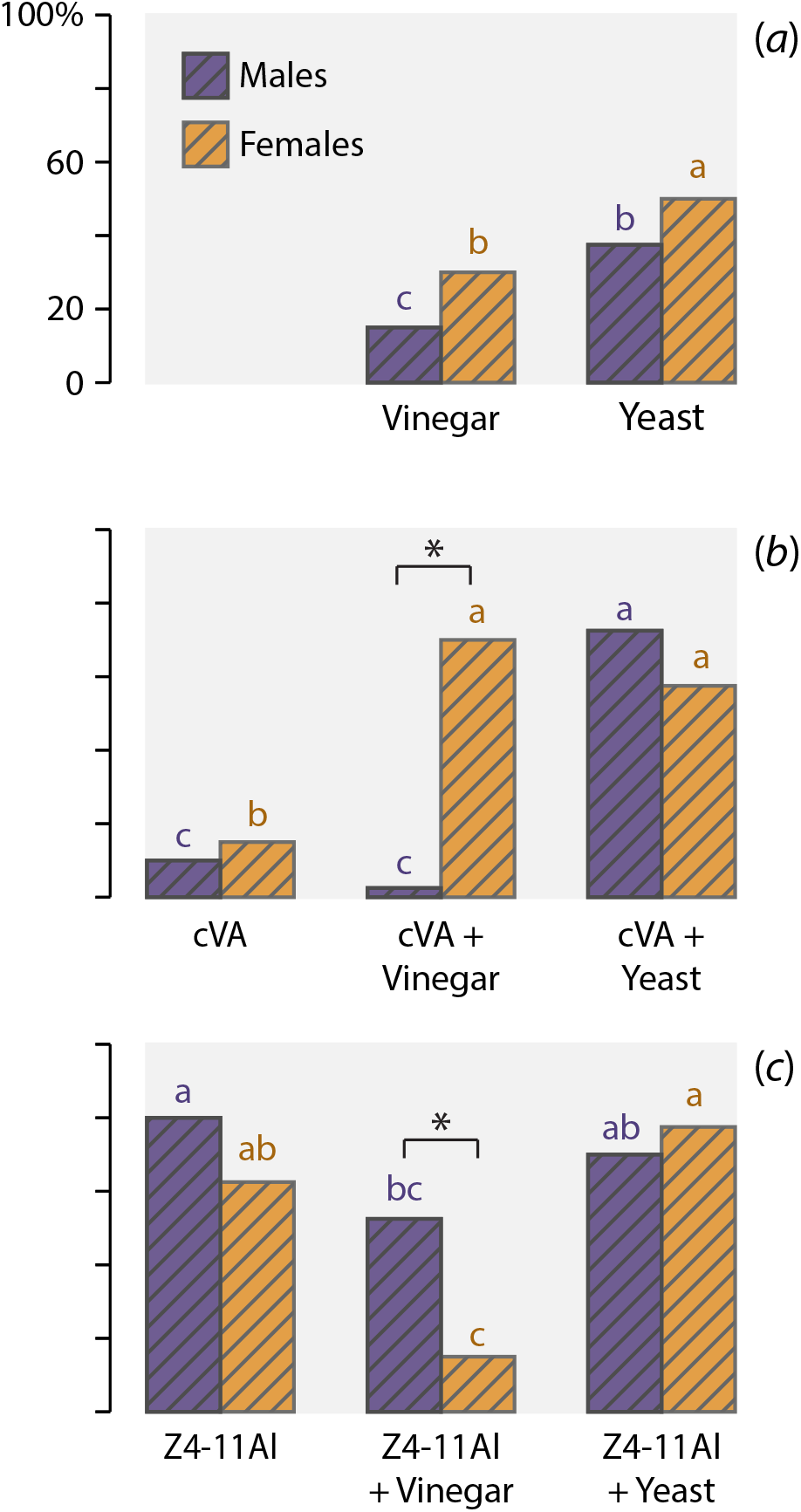
Odour-mediated upwind flight attraction of fruit fly *Drosophila melanogaster* males (blue bars) and females (ochre bars) to (a) vinegar and yeast headspace, (b) 10 ng/min male sex pheromone cVA, (c) 10 ng/min female sex pheromone *Z*4-11Al, alone and in blends with vinegar or yeast headspace, respectively. Letters of the corresponding colours show differences between treatments using Wald pairwise comparisons. Asterisks show treatments with significant differences between sexes (*n*=40).

Male pheromone cVA alone attracted only few flies, while a blend of cVA and vinegar was highly attractive to females, not to males (Figure 1b). In comparison, both sexes responded to a blend of cVA and yeast headspace.

Female pheromone *Z*4-11Al alone attracted males and females. Blending vinegar with *Z*4-11Al reduced attraction in both sexes, but significantly more males than females responded. And, a blend of yeast headspace and *Z*4-11Al attracted as many flies as *Z*4-11Al alone (Figure 1c).

Yeast headspace by itself attracted more males and females than vinegar headspace (Figure 1a). Females attraction to yeast headspace (Figure 1a) and to the blends of yeast headspace and cVA and *Z*4-11Al, respectively, was not significantly different (Figures 1b,c).

Taken together, only few flies responded to the male pheromone cVA, while the female pheromone *Z*4-11Al strongly attracted both sexes. Blending vinegar with cVA and *Z*4-11Al, respectively, produced an inverse response pattern in male and female flies. Our results show further that vinegar, a frequently used food odorant source for *Drosophila,* is an inferior attractant in comparison with yeast aroma (Figure 1).

### Male courtship in response to *Z*4-11Al

Male-produced cVA mediates aggregation, suppresses male-to-male courtship, and males learn to avoid mated females tainted with cVA (Ejima et al. 2007, Kurtovic et al. 2007, Keleman et al. 2012). Since female-produced *Z*4-11Al elicits male flight attraction (Figure 1c), we asked whether *Z*4-11Al also has an effect on courtship, following landing at females. Males were tested with decapitated target flies, laced with blank solvent or synthetic test chemicals.

Compared with solvent control, *Z*4-11Al elicited a significant response in experienced males that had mated earlier, and not in naive, unmated males. Males scored showed vigorous courtship including attempted copulation (Figure 2a). More experienced males courted decapitated females than males, plausibly because cVA, present on the cuticula of male flies, has an antagonistic effect on experienced males. That a few males nonetheless courted decapitated males further confirms a role of *Z*4-11Al in male courtship (Figure 2a). Tests with Canton-S test males corroborated results obtained with males of the Dalby strain (Figure 2b).

**Figure 2.**
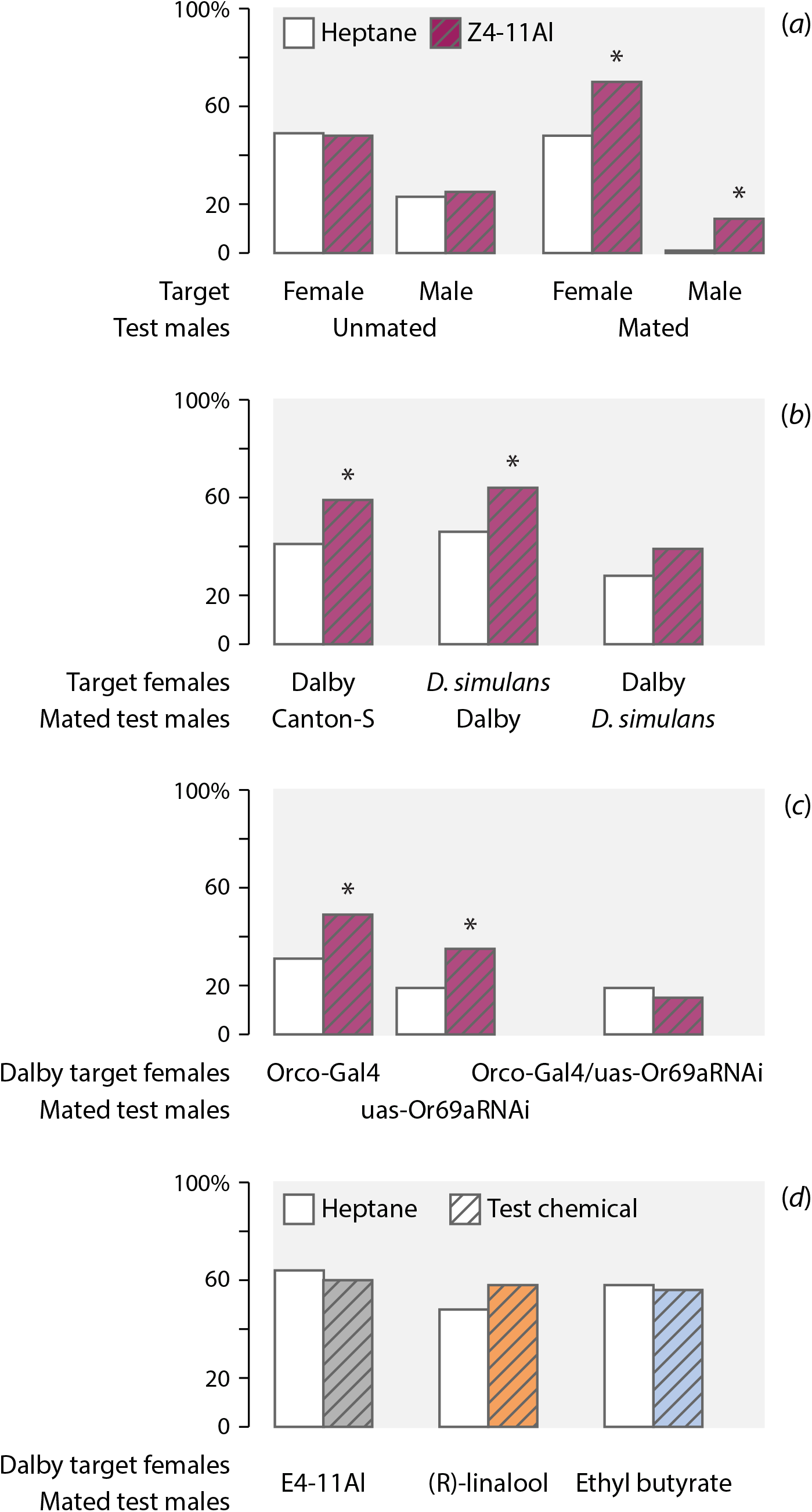
Effect of *Z*4-11Al on courtship in *D. melanogaster* males. Decapitated target flies were painted with 1 ng test compound or heptane (solvent control). The number of courting and non-courting males in each test was compared using a chi-square test, asterisks show significant differences. (a) Proportion of unmated or mated *D. melanogaster* (Dalby) test males courting unmated decapitated female or male target flies treated with 1 ng *Z*4-11Al or heptane (solvent control) (*n*=80, *P*=0.006 and *P*=0.007). (b) Proportion of mated Canton-S and Dalby strain *D. melanogaster* males and *D*. *simulans* males, courting decapitated Dalby or *D*. *simulans* females, treated with 1 ng *Z*4-11Al or heptane (*n*=80, *P*=0.009 and *P*=0.039). (c) Effect of RNA interference, in Orco-Gal4/uas-Or69aRNAi mated males, courting Dalby target females, treated with 1 ng *Z*4-11Al or heptane. Parental lines, Orco-Gal4 and uas-Or69aRNAi, show significant courtship behaviour (*n*=80, *P*=0.036 and *P*=0.032). (d) Proportion of mated Dalby males courting decapitated Dalby females, treated with *E*4-11Al, (*R*)-linalool or ethyl butyrate (*n*=80).

Importantly, males also courted decapitated *D*. *simulans* target females, laced with synthetic *Z*4-11Al (Figure 2b). This experiment rules out a contributing role of the cuticular hydrocarbon pheromone 7,11-HD, which is specific for *D*. *melanogaster* and not found in *D*. *simulans* females (Billeter et al. 2009), and underlines that *Z*4-11Al elicits male courtship, in the absence of 7,11-HD.

On the other hand, *D*. *simulans* males did not respond in a significant manner to Dalby females painted with *Z*4-11Al (Figure 2b). This was expected, since *D*. *simulans* males are not attracted to *Z*4-11Al (Lebreton et al. 2017) and since *D*. *simulans* females do not produce 7,11HD, which is the precursor for *Z*4-11Al (Billeter et al. 2009, Lebreton et al. 2017).

Moreover, we used a RNAi fly line to corroborate that Or69a encodes *Z*4-11Al-mediated courtship. Significantly fewer males responded when olfactory sensory neurons (OSNs) expressing Or69a were disrupted (Figure 2c).

Finally, we tested three non-pheromonal chemicals: *E*4-11Al, the geometric isomer of *Z*4-11Al, which is perceived via the Or69aB variant; (*R*)-linalool, a food compound, which is the main ligand of the other variant, Or69aA; and ethyl butyrate, which is a ligand for Or22a (DM2 glomerulus in the AL), for Or43b (VM2 glomerulus) and also for Or85a (DM5 glomerulus; see also Figure 3). None of these compounds elicited a significant courtship response in experienced males (Figure 2d).

**Figure 3.**
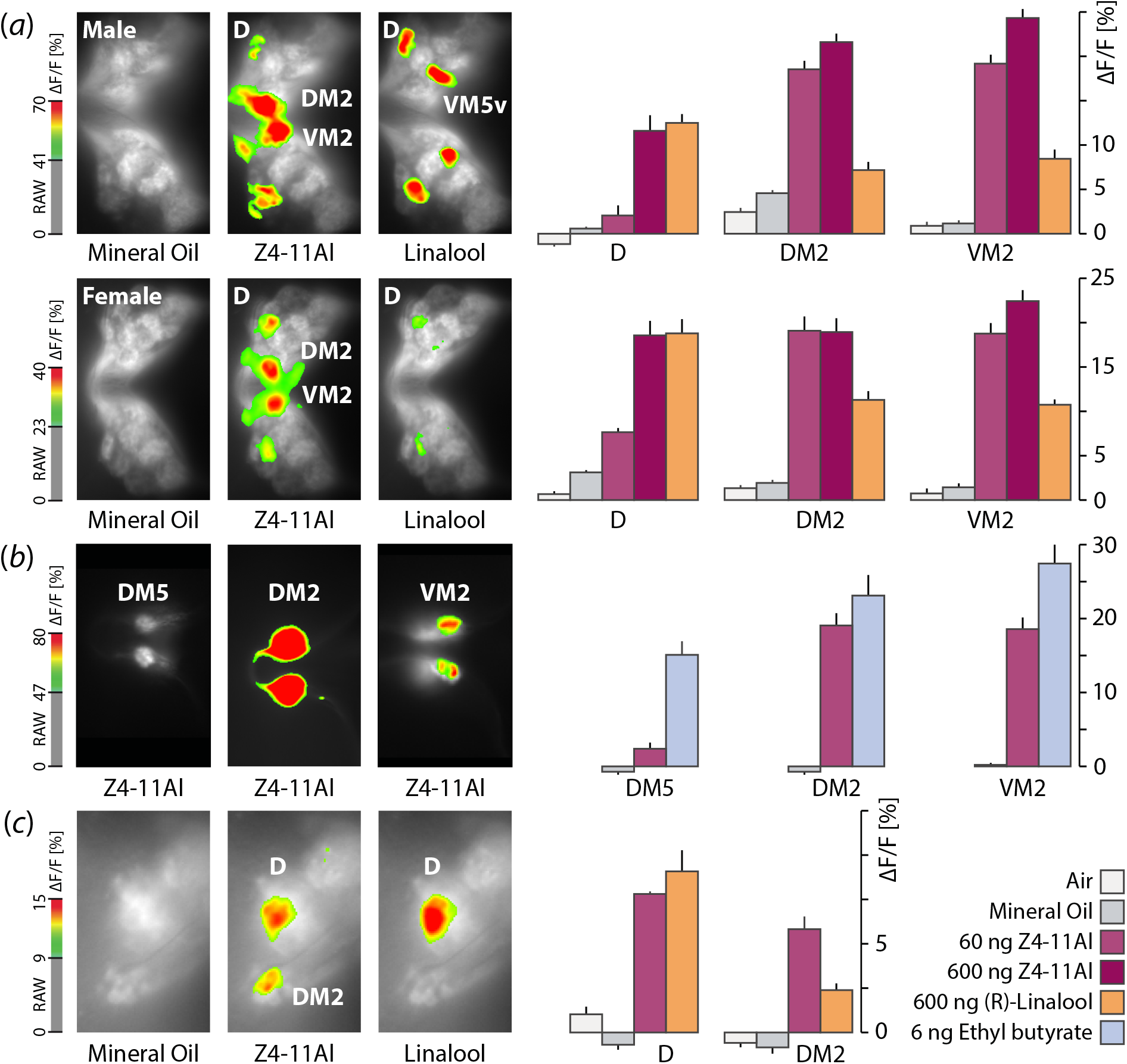
Calcium imaging responses in the antennal lobe (AL). Colours show the median normalized calcium activity (Δ F/F [%]) in response to controls and odor applications, according to the colour bar on the left (*n*=10; mean±SD). (a) Calcium activity in Orco-Gal4 males (top) and females (below) in response to control (air, mineral oil), to 60 and 600 ng *Z*4-11Al, and to 600 ng (*R*)-linalool. *Z*4-11Al and (*R*)-linalool are key ligands for the D glomerulus (Lebreton et al. 2017). (b) Responses of Or22a-Gal4 males to *Z*4-11Al in the DM2 glomerulus and of Or43b- Gal4 males in the DM5 and VM2 glomeruli; ethyl butyrate is a diagnostic stimulus for these glomeruli. (c) Projection neuron (PN) responses in GH146-Gal4 males to *Z*4-11Al and (*R*)-linalool in D and DM2 glomeruli.

### Activation of antennal lobe glomeruli in response to *Z*4-11Al

The D glomerulus in the antennal lobe (AL) collects input from ab9A olfactory sensory neurons (OSNs) that co-express Or69aA and Or69aB, the two isoforms of Or69a (Couto et al. 2005). Single sensilllum recordings (SSR) have further shown that the food odour (*R*)-linalool and female pheromone *Z*4-11Al are the respective key ligands for Or69aA and Or69aB (Lebreton et al. 2017).

We used calcium imaging of the AL to first confirm the role of the Or69a channel including the D glomerulus in the perception of *Z*4-11Al. Moreover, we compared the AL response to the key ligands (*R*)-linalool and *Z*4-11Al. Both compounds elicit upwind flight (Lebreton et al. 2017), but only *Z*4-11Al participates in male courtship, which goes to show that the flies discriminate between these two compounds (Figure 2).

In males and females, *Z*4-11Al activated the D glomerulus, as expected, but also DM2 and VM2 (Figure 3a). These broadly-tuned glomeruli have also been shown to respond to a mix of cVA and vinegar (Lebreton et al. 2015). However, neither ab3A OSNs (expressing Or22a, connecting to DM2), nor ab8A (expressing Or43b, connecting to VM2; Grabe et al. 2015, Münch and Galizia 2016) responded to *Z*4-11Al, according to SSR (Lebreton et al. 2017). Simultaneous activation of DM2 and VM2 glomeruli by *Z*4***-*** 11Al might accordingly reflect interglomerular communication in the AL, via local neurons (Wilson 2013).

(*R*)-linalool, on the other hand, is also a ligand for ab7A OSNs expressing Or98a, which accounts for additional activation of the VM5v glomerulus (Couto et al. 2005, Münch and Galizia 2016). In comparison, activation of DM2 and VM2 glomeruli by (*R*)-linalool was much lower compared to *Z*4-11Al (Figure 3a).

Activation of the DM2 and VM2 glomeruli by *Z*4-11Al was substantiated by imaging Or22a-Gal4 males and Or43b-Gal4 males, respectively. Ethyl butyrate is a diagnostic stimulus for Or22a, Or43b and Or85a (DM2, VM2 and DM5 glomeruli) (Figure 3b; Grabe et al. 2015, Münch and Galizia 2016).

Imaging of the glomerular activation pattern of projection neurons (PNs), connecting the AL to the lateral horn (LH), where behavioural responses are generated, corroborated activity of *Z*4-11Al in the D and DM2 glomeruli (Figure 3c). A possible response of VM5v to linalool does not show, since GH146-Gal4 does not label this glomerulus (Grabe et al. 2015).

## Discussion

Volatile chemicals that are cognate ligands for odorant receptors (ORs) are selectively perceived against noisy backgrounds and they capacitate behavioural decisions at a distance, since they deliver reliable information about the source, including identity and physiological state of the emitter, in the case of pheromones.

The female pheromone *Z*4-11Al attracts naive males and females by flight (Figure 1). At close range, *Z*4-11Al also elicits courtship, but only in experienced, previously mated males (Figure 2). A response in naive males is not expected, since neurons mediating perception of *Z*4-11Al are not part of the sexually dimorphic fruitless circuitry that elicits innate male courtship (Manoli et al. 2005, Stockinger et al. 2005).

Importantly, males respond also to decapitated *D*. *simulans* females laced with *Z*4-11Al. This shows that a response to *Z*4-11Al does not require presence of the close-range CHC pheromone 7,11-HD, since 7,11-HD is not produced by *D*. *simulans* (Billeter et al. 2009, Lebreton et al. 2017, Billeter and Wolfner 2018, Sato and Yamamoto 2020).

### Courtship discrimination between Or69a ligands

Female pheromone Z4-11Al and the food odorant (*R*)-linalool are main ligands for Or69aB and Or69aA, which are co-expressed in ab9A OSNs, branching to the D glomerulus in the AL (Couto et al. 2005, Lebreton et al. 2017).

Whereas both compounds elicit innate flight attraction (Lebreton et al. 2017), males distinguish between Z4-11Al and (*R*)-linalool during courtship (Figure 2a,d), probably because males learn to associate Z4-11Al with females during mating. Courtship learning confirms production of Z4-11Al by females. The question arises whether males would even learn to associate (*R*)-linalool or other food-related compounds with females.

A differential response to Z4-11Al and (*R*)-linalool is reflected by their respective activation patterns in the AL (Figure 3a). (*R*)-linalool is also a ligand for Or98a, which accounts for stimulation of the VM5v glomerulus (Couto et al. 2005, Münch and Galizia 2016)

In contrast, direct activation of DM2 and VM2 by Z4-11Al is not supported by SSR data (Lebreton et al. 2017) and lateral activation by local AL neurons is a possible explanation, instead (Wilson 2013). DM2 and VM2 are respond broadly, also to blends of vinegar and cVA (Lebreton et al. 2015).

Finally, we do not know whether the olfactory input of Or69aA and Or69aB through ab9A is entirely equivalent. OSN spiking trains generated by Or69aA and Or69aB may deliver non-congruent messages that may be distinguishable at the AL level. This is reminiscent of Or85e and Or33c which are coexpressed in ab3A OSNs, where it is yet unclear whether different response profiles and spiking patterns enable behavioural discrimination between the respective Or ligands (Goldman et al. 2005).

### Roles of cVA and Z4-11Al in upwind flight and courtship

Male courtship in *D*. *melanogaster* has been anatomically and physiologically dissected at a neuronal circuit level, with a particular emphasis on the sex-dimorphic behavioural effect of cVA. A central node is the sexually dimorphic P1 interneuron cluster that integrates input from all sensory modalities to activate male courtship, and that is regulated by social experience. Male-produced pheromone cVA provides antagonistic olfactory input to the P1 node, preventing courtship of recently mated females perfumed with male cVA (Manoli et al. 2005, Stockinger et al. 2005, Kimura et al. 2008, Ruta et al. 2010, Kohl et al. 2013, Yamamoto and Koganezawa 2013, Clowney et al. 2015, Kohl et al. 2015, Auer & Benton 2016).

cVA does not elicit long-range flight by itself, only in combination with fly food (Figure 1; Bartelt et al. 1985, Lebreton et al. 2015) and is shared by many other *Drosophila,* including the sibling species *D*. *simulans* (Schaner et al. 1987, El-Sayed 2020). Females, on the other hand, produce the species-specific CHC 7,11-HD that feeds gustatory input into P1 to activate male courtship and to achieve reproductive isolation towards the sibling species *D*. *simulans* (Billeter et al. 2009, Thistle et al. 2012, Toda et al. 2012, Seeholzer et al. 2018). What is yet unclear is whether P1 also receives excitatory olfactory input.

Does Z4-11Al feed into P1? Z4-11Al encodes species-specificity, mediates long-range attraction (Figure 1; Lebreton et al. 2017) and contributes to courtship, independently of 7,11-HD (Figure 2a,b). Z4-11Al is perceived prior to and during upwind flight, and since females release Z4-11Al, it follows that stimulation is sustained at close-range. Courtship and upwind flight are not isolated episodes, they are inextricably interconnected parts of a behavioural sequence that culminates in copulation.

### A behavioural paradigm for bioactive odorant identification

An unambigious, stereotypical behavioural response is substrate for investigations of the genetic basis, neural circuitry and physiology of mate communication in *D*. *melanogaster.* We here show that upwind flight attraction is a behavioural paradigm that is suitable to extend investigations of reproductive behaviour to include long-range communication, and even decision-making upon odorant stimulation.

Comparison of pheromone blend attraction with vinegar and yeast shows that yeast overrides differences between pheromones (Figure 1). Vinegar, derived from acetic acid bacteria fermentation (Lynch et al. 2019), is a widely used standard food attractant for *D. melanogaster*, although live yeast aroma is much richer in composition (Callejón et al. 2009, Chinnici et al. 2009, Becher et al. 2010, Ljunggren et al. 2019). Yeast growing on ripe fruit is also a biologically more relevant attractant, since it attracts flies for oviposition and since yeast is a sufficient substrate for larval development (Becher et al. 2012, Grangeteau et al. 2018, Quan and Eisen 2018, Murgier et al. 2019).

Reproductive behaviour in *D*. *melanogaster* is mediated by an ensemble of pheromones and food odorants, emanating from food substrates and from aggregating or mating flies. Our knowledge of *D*. *melanogaster* pheromone chemistry is fairly comprehensive, compared to behaviourally salient food odorants, owing to the extraordinary complexity of microbial headspace.

Flies recognize food sources despite very considerable, inherent variation in the bouquets emanating from fermenting fruit, according to fruit substrates, microbial community composition and fermentation state. Towards an understanding of how neural representations of such complex and variable odours are generated (Endo et al. 2020) and how habitat cues are integrated with pheromonal signals we next need to identify the key odorants in microbial food headspace. An upwind flight bioassay is suitable to describe the key bioactive chemical constitutents that encode reliable recognition and evaluation of food odour objects and trigger the decision to engage in orientation flights.

## Acknowledgements

We thank two anonymous reviewers for constructive criticism.

## Data availability

Data will be made avaliable at Dryad.

